# Changes of sensory and emotional tactile dimensions with age in self-touch via the application of creams

**DOI:** 10.64898/2026.09.04.748740

**Authors:** Maria Rosa Bufo, Philippe Bastien, Mariama Dione, Aurélie Coubart, Jean-Marc Aimonetti, Roland Jourdain, Rochelle Ackerley

**Affiliations:** Aix Marseille Univ, CNRS, CRPN (Centre de Recherche en Psychologie et Neurosciences – UMR 7077), Marseille, France; L’Oréal Research & Innovation, Aulnay-sous-Bois, France; L’Oréal Research & Innovation, Chevilly-Larue, France

**Keywords:** affective touch, aging, touch perception, statistical models, humans, hairy skin, finger, glabrous skin

## Abstract

Touch moderates how we interact with objects, others, and ourselves, where self-touch is particularly important in self-care, although fewer studies have investigated its perception, especially with age. Here, women aged 20-28 and 65-75 years rated how self-applied creams felt on the cheek and forearm, using sensory and emotional descriptors in the Touch Perception Task. Factor analyses uncovered higher tactile dimensions, including Texture, Moisture, Positive Affect, and Negative Affect. Sensory dimensions were more intense in the young group, whereas Positive Affect was stronger in the older group. We then used network and clustering analyses to map how the descriptors interacted. Young participants showed significantly higher connectivity between descriptors, whereas older participants displayed fewer connections, but many became mixed sensory-emotional pairings. Clustering analyses revealed further changes in the tactile groupings with age. We conclude that aging reshapes touch into an emotionally-rich experience, which impacts how interact with ourselves and others.

## Introduction

Touch is a proximal sense that infers contact with the body, where tactile interactions can have profound effects, shaping our experiences and reactions to them. Touch has often been separated into sensory-discriminative (e.g. spatial acuity, touch detection) and emotional-affective (e.g. pleasant, arousing) dimensions [1,2], but nearly all tactile experiences inherently combine these [3]. Although we require the sensory side of touch to interact with objects efficiently, the evoked emotional state drives our behavior in daily life, for example, whether we choose to repeat an interaction [4]. Much work has demonstrated the importance of touch in inter-personal interactions and in social relationships [5,6], including in situations like grooming and self-care [7]. Further, self-touch is important, yet a comparatively less-investigated form of touch, that also has strong implications in self-care and maintaining our sense of self [8]. To this end, the hands are fundamental in self-touch, whether it be hand-to-hand contact in bimanual tasks or hand-to-body contact, although certain parts of the body are more easily reached (e.g. face, arms) than other parts (e.g. back, feet).

Skin can be classified as glabrous or hairy, containing different afferents at different densities [5,9]. All skin contains fast-conducting, low-threshold Aβ mechanoreceptive afferents that are linked to discriminative-sensory touch capacities, where their density is maximal on the hands and face, giving very fine tactile acuity at these areas [10,11]. Skin also contains a further type of low-threshold mechanoreceptor, which is unmyelinated and slowly-conducting, namely C-tactile (CT) afferents, also known a C low-threshold mechanoreceptors (CLTMRs). These are abundant in the hairy skin of the arm [12] and have also been found on the face [13], whereas they are rare on the glabrous skin of the hands [14] and relatively rare on the leg [15]. Although Aβ mechanoreceptor input can be used centrally to provide emotional tactile value [3], CT afferents have been implicated directly in signaling positive affective touch, where their intensity of firing correlates with pleasantness ratings [16,17]. It is important to consider the skin being touched, to understand better how tactile experiences are interpreted, where the upper half of the body is more sensitive to touch [10,11,18] and temperature [19,20], as well as being more involved in the embodiment of emotions [21].

Self-touch influences our body image and sense of self throughout our lives, starting prenatally [22], which aids us in distinguishing ourselves from others and objects. Self-touch is particularly intriguing due to the suppression of active movements towards the body, which is why we find it difficult to tickle ourselves [23,24]. However, the suppression of sensory feedback from active movements in self-touch can be regulated, where the extent signal cancellation may depend on other factors, such as attention towards the skin being stimulated [25,26]. Active self-touch necessitates attention, such as in grooming and bimanual hand gestures, where the hand is key, providing both discriminative and affective information. Although Aβ mechanoreceptor density is high on the hands and CT density very low, the hands give a relatively similar contribution to the perception of pleasant touch, even when compared to CT-dense regions like the forearm [27]. Thus, when we touch our own skin, such as in a self-care routine, the sensation of the hand moving over the skin produces a complex interaction of tactile input from both skin areas, as well as the additional effects of anything we have on the skin, such as in applying creams to the skin or through clothes. The inter-dependence between active hand-face self-touch was recently demonstrated where having dry hands made the facial skin feel dry, even when it was not [28] and that tactile learning can transfer from the hand to the face [29]. Touch to the forearms also forms part of many tactile expressions in humans [30], where the whole upper body is highly implicated in the experience of emotions [21].

Touch also changes throughout our lives, where self-touch plays a central role in establishing and maintaining the sense of self and in self-care. From adulthood, there is a 5-8% decrease in the number of mechanoreceptive afferents per decade [9] and its impact is seen in the diminution in tactile sensitivity with age, although this mostly manifests on the glabrous skin of the hands and feet [9,31]. Conversely, tactile pleasantness has been found to increase with age [28,32,33], possibly due to an increase in the appreciation of touch, a change in the balance of mechanoreceptive afferent input, and/or other cognitive or lifestyle differences. Therefore, even if the hairy skin is less affected by age, self-touch using the hands will inevitably change over the lifespan, due to the large impact of aging on the glabrous hand skin.

Various methods can be used to measure tactile perception, including standard tests of touch sensitivity and spatial discrimination [34], as well as perceptual rating scales (e.g. roughness, pleasantness) [35,36]. However, these measures are typically designed to capture one component of tactile perception, yet touch is multi-dimensional. The Touch Perception Task was formulated to capture the wider dimensions underlying touch, which consists of rating 26 sensory and 14 emotional somatosensory descriptors [2]. It has been translated into other languages and the approach has been adapted in various ways [1,37–41]. These studies have shown that sensory-discriminative aspects of touch can be captured in various higher-level concepts, including roughness, slip, firmness, pile, texture, moisture, fluidity, and granularity. Emotional-affective aspects are also found, but tend to be fewer, where a dimension relating to positive affect is often found, as well as a dimension corresponding to arousal, and sometimes negative affect. This task has been used to investigate touch perception, successfully capturing material properties [1,2,37,39]. Further, differences between body sites can be found, especially relating to emotional dimensions, and between the mode of touch (e.g. active, passive), where passively-applied touch tends to feel more intense [1,2,38].

We presently investigated tactile dimensions using the Touch Perception Task, through the self-application of a cream onto the skin of the cheek and forearm, to explore how self-touch is perceived in a self-care, considering the importance of the hands, face, and arms. Our aim was not to determine the sensory and emotional feel of creams *per se* [37–39], as this was assessed in a previous, related study [28], but to investigate how self-touch is perceived during their application and whether this changed with age, predicting that emotional dimensions would dominate in an older group, compared to younger participants. We had a further objective to apply advanced analytical methods to the Touch Perception Task, to not only uncover different tactile dimensions using conventional factor analyses, but to take this further to map how these dimensions evolve with each other.

## Methods

Two age groups were recruited for the present work with inclusion criteria being either a self-described healthy woman between 20-30 or 65-75 years’ old. Exclusion criteria were pregnancy or lactation within the past 6 months, as well as having a history of neurological, psychiatric, and/or dermatological issues, or clinically significant peripheral neuropathy, such as diabetes. For the young group, 22 participants (median age: 24 years, range: 20-28 years) completed the study. For the older group, 22 participants were recruited, but one was found to have various issues and was removed from the study, thus, 21 participants (median age: 70 years, range: 65-75 years) were included. The study was approved by the ethical committee Est-III in France (no. 21.02.02.10694) and was performed in accordance with the Declaration of Helsinki, apart from pre-registration. Written informed consent was obtained, and participants were paid for their time. The present investigation was part of a larger experimental study on tactile sensations and aging [28].

### Experimental procedures

Each participant came for two separate experimental sessions, where one of two similar emulsion creams was tested (Cream A and Cream B) and the order of the cream was randomized between participants/sessions (see [28] for further details). The products were in identical, unbranded pots and had the same light perfume. The tested creams were rather similar emulsions and the previous study found no significant differences between them on a number of variables [28], thus they are included here as a repetition of the task. Participants sat at a table in a quiet room, with a constant temperature of ∼22°C. They first watched a video describing how to rub in the cream, via self-touch, using gentle circular motions. The experimenter applied 0.05 ml of cream using a micropipette to give four drops either on the arm, or on the right index finger of the participant (on the distal phalanx). The participant then rubbed the cream into either the left arm or the left cheek, respectively, using their right index finger over an area of ∼10 cm^2^. After this application the participant answered the Touch Perception Task about the sensations. The order of body site touch was randomized between participants.

### Touch Perception Task

The Touch Perception Task was used to examine differences in sensory (discriminative) and emotional (affective) aspects of touch at different skin sites. For this, a total of 26 sensory and 14 emotional somatosensory descriptors are used [1,2]. The original task was in English and was translated into French for the present experiment (Table 1). An initial translation was completed by three bilingual people and this was back-translated by a further three different people. The words were then matched and verified. After the participants had applied the cream to the arm or cheek, the touch descriptors were given on a piece of paper and the participant had to rate how descriptive each word was by crossing one descriptor box, composing of “Not descriptive”, “Slightly descriptive”, “Moderately descriptive”, “Highly descriptive” and “Very highly descriptive”. These descriptors were coded onto an ordinal scale of 0-4, respectively. Each participant completed this for the two creams at the two sites.

**Table 1.**
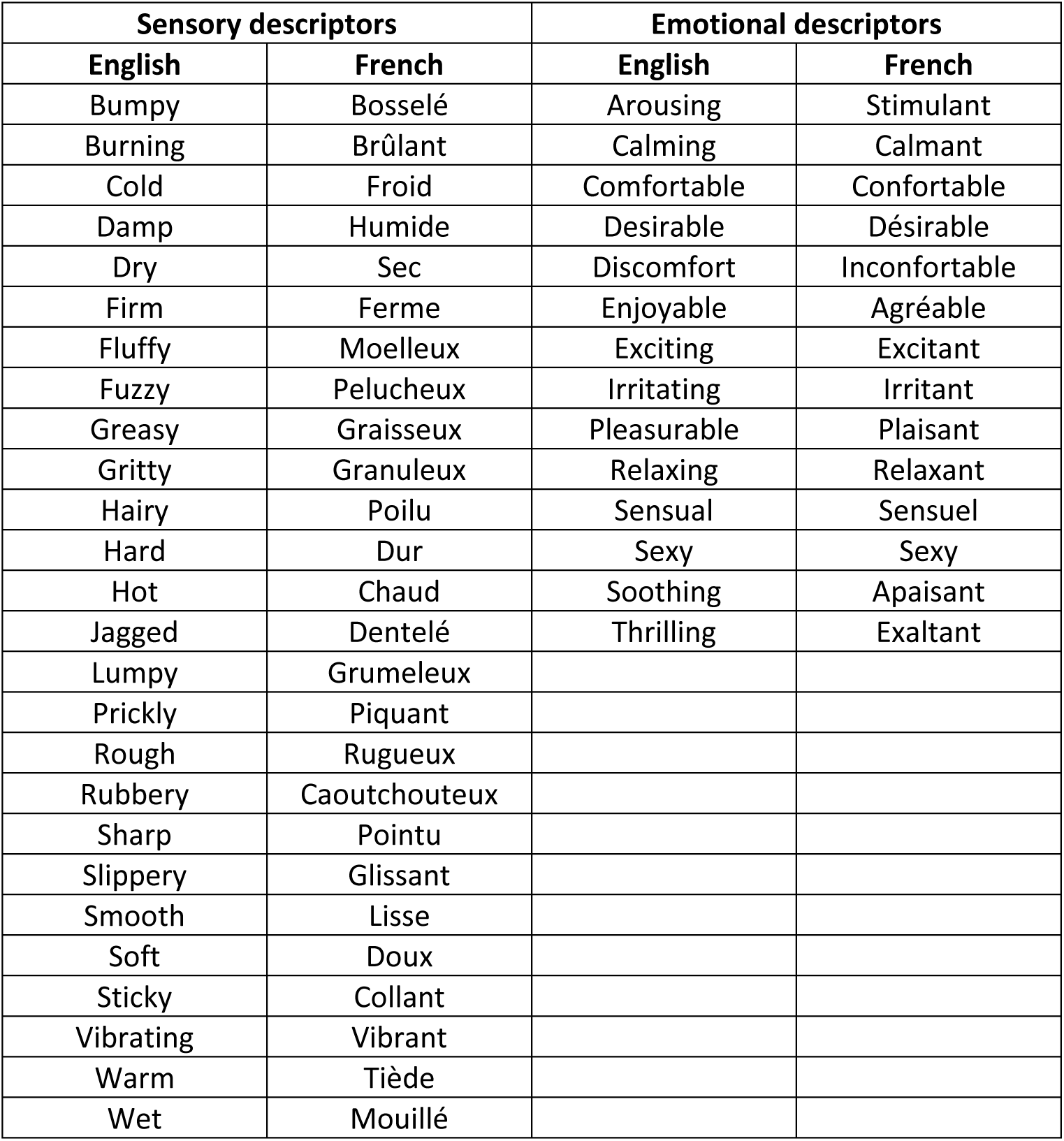
Translation from English to French of the Touch Perception Task.

| Sensory descriptors |  | Emotional descriptors |  |
| --- | --- | --- | --- |
| English | French | English | French |
| Bumpy | Bosselé | Arousing | Stimulant |
| Burning | Brûlant | Calming | Calmant |
| Cold | Froid | Comfortable | Confortable |
| Damp | Humide | Desirable | Désirable |
| Dry | Sec | Discomfort | Inconfortable |
| Firm | Ferme | Enjoyable | Agréable |
| Fluffy | Moelleux | Exciting | Excitant |
| Fuzzy | Pelucheux | Irritating | Irritant |
| Greasy | Graisseux | Pleasurable | Plaisant |
| Gritty | Granuleux | Relaxing | Relaxant |
| Hairy | Poilu | Sensual | Sensuel |
| Hard | Dur | Sexy | Sexy |
| Hot | Chaud | Soothing | Apaisant |
| Jagged | Dentelé | Thrilling | Exaltant |
| Lumpy | Grumeleux |  |  |
| Prickly | Piquant |  |  |
| Rough | Rugueux |  |  |
| Rubbery | Caoutchouteux |  |  |
| Sharp | Pointu |  |  |
| Slippery | Glissant |  |  |
| Smooth | Lisse |  |  |
| Soft | Doux |  |  |
| Sticky | Collant |  |  |
| Vibrating | Vibrant |  |  |
| Warm | Tiède |  |  |
| Wet | Mouillé |  |  |

### Statistics

The data gained from the touch perception task were ordinal. To analyze the touch descriptors, two separate analyses were undertaken using R Studio (version 2024.09.0) and Python (version 3.11). We first explored the dataset and we observed descriptor that had unbalanced modality scores (Supplementary Figure 1), which was observed for both young and older groups. This prevented us from conducting robust statistical analyses. Therefore, to address this issue, we recoded the dataset to increase variability, while preserving interpretability. Specifically, items scored as zero (Not descriptive) were retained as such. Items originally scored as one or two (Slightly descriptive and Moderately descriptive, respectively) were recoded as one, and scores of three or four (Highly descriptive and Very highly descriptive, respectively) were recoded as two. This resulted in a simplified scoring system ranging from 0 (not at all descriptive) to 2 (very descriptive), thereby improving the suitability of the data for network analysis to make predictions from the data.

As outlined in detail below, we conducted three main types of analysis to model the relationship between sensory and emotional aspects of touch, including how these change with age and body site, and to explore whether predictions can be made between these. In the first analysis, we performed exploratory factor analyses to explore sensory and emotional descriptors, with ANOVAs to look at age and body site effects. Normality was assessed using the Shapiro–Wilk test and homogeneity with Levene’s test. These linear analyses have been classically carried out in the literature on touch descriptors [1,2,37], thus, for comparability with previous work, we completed this first. In the second part, we performed a network analysis to investigate partial correlation between descriptors, to further investigate the interdependencies between sensory and emotional descriptors. In the third approach, we explored the sensory and emotional factors together further, where we conducted non-linear clustering analyses to investigate relationships between grouping descriptors, i.e. the way in which they clustered.

### Factor analysis

In the first step, we performed three exploratory factor analyses. Firstly, we conducted two factor analyses to explore sensory and emotional descriptors separately, as per the literature [1,2,37], and we then explored them together, as we had no premise that these should be distinct, making three different analyses (sensory, emotional, sensory + emotional). For ease of comparison, we labeled the sensory descriptors in blue and the emotional descriptors in red. We used a polychoric correlation matrix for our data, which is better suited for ordinal data. For comparability in the factor analyses, we used the classic approach in the literature [1,2]. In brief, Kaiser-Meyer-Olkin measures of sampling were gained for factor analysis to verify that a sufficient sample size was used. This measure was 0.79 for the sensory factor analysis, 0.87 for the emotional factor analysis, and 0.80 for the combined sensory + emotional factor analysis, which are all well above the 0.7 acceptability limit [42]. Bartlett’s test was used to check the significance of the correlation matrix of factors and was found to be non-significant in all cases, indicating that the descriptors cluster into independent factors. Significant factors were determined by having eigenvalues of >1 and accounting for >5% of the variance. We kept descriptors which had loadings of >0.3. The names of the factors were chosen according to the descriptors that loaded with high values on that factor from the regression matrix of factor scores, which was performed with orthogonal varimax rotation output. Statistical analyses were then conducted on the regression matrix score outputs from all participants, where an ANOVA was realized for each factor, testing age group (levels: Young, Older) and body site (levels: Arm, Cheek) as conditions. Significance was set at p < 0.05 and values are expressed in the figures as means ± 95% confidence intervals.

### Network analysis

We performed a second analysis: a network analysis based on graphical lasso [43], to investigate the relationships and interdependencies between sensory and emotional descriptors. Prior to this, polychoric correlations were performed to fit the ordinal category of our touch descriptors, which is necessary to derive assumptions about the results. To identify the nodes that likely measured the same underlying construct (i.e. that were colinear), the Goldbricker wraps package in R Studio was used, which compares every possible combination of correlations in a psychometric network. It calculates the proportion of correlations that are significantly different for each different pair of nodes [44]. We used 0.25 as a threshold and all pairs of nodes that fell below this threshold were returned as ‘bad pairs’ and removed. Only two pairs were removed, namely Sexy-Exciting with a score of 0.13 and Sensual-Desirable with a score of 0.24. Subsequently, we used graphical lasso models to estimate the network. Sparsity parameters were also extracted, which represent the proportion of edges that are non-zero (i.e. have a connection) compared to the total number of possible edges, where sparsity values between 0.05-0.3 are common [45].

In order to verify the network structure, we extracted network parameters, namely node strength and node betweenness, with node strength the sum of the weights of all direct connections to a node, while node betweenness measures how often a node lies on the shortest path between other nodes in the network. Network stability was evaluated through centrality stability coefficients and bootstrapped edge-weight confidence intervals. Networks with a centrality measure coefficient above 0.5 for nodes and narrow confidence intervals for edges are considered stable [46–48]. Thus, we used a tuning parameter of 0.5 and non-parametric bootstrap reiteration with 2000 resamples (Epskamp et al., 2017, 2018; Friedman et al., 1993; Friedman et al., 2008).

To compare between conditions, we performed network comparison tests for independent groups between the two networks representing the different age groups (young and older participants) and for dependent groups between the two networks representing different body sites (cheek and arm). Network comparison tests are used to compare networks on various levels, such as the overall structure and individual edges [48]. We report a test statistic (M) for overall network structure invariance, which informs us of whether the two networks overall structures were significantly different. Clustering coefficient testing was used to understand the extent to which nodes clustered together, which we calculated globally (i.e. for the entire network). We used the non-parametric permutation test (also known as a randomization test), to assess the statistical significance of differences between clustering coefficients. We used the false discovery rate method to adjust the p-values to control for multiple comparisons.

### Clustering analysis

In the third analysis, we used a non-linear approach to cluster the data and see how the descriptors grouped together, to add predictive value. This procedure was applied to both the young and older groups to visualize how dimensions of the tactile experience of touch varied across age groups. Body site differences were not considered due to non-significant results for this in the network analyses. We applied the t-distributed stochastic neighbor embedding (t-SNE) algorithm for dimensionality reduction, to reduce the high-dimensional sensory and emotional data into a lower-dimensional space, making it manageable for classification. This was done using a perplexity value of 4 and an early exaggeration of 12. Perplexity controls the balance between local and global aspects of the data, while early exaggeration helps separate clusters early in the learning process. The t-SNE results were plotted to visualize the emotional and sensorial variables in two dimensions, where similar variables appear closer together. Subsequently, KMeans clustering was applied to the t-SNE results to group the emotional and sensorial variables into clusters. The KMeans algorithm identified patterns within the data by grouping similar variables based on their proximity in the t-SNE space and each cluster was labeled with descriptive names reflecting the underlying characteristics of the group.

## Results

### Factor analyses

In three separate factor analyses, sensory and emotional aspects were explored to assess the effects of age (young, older) and body site (cheek, forearm) on each domain. For the sensory factor analysis, four dimensions were found: Texture, Moisture, Friction, and Temperature. Table 2 shows the variance explained per factor and the regression loadings. Using ANOVA, we found a significant main effect of age group for Texture (F_(1, 168)_ = 6.19, p = 0.014), Moisture (F_(1, 168)_ = 19.75, p < 0.001), and Temperature (F_(1, 168)_ = 16.54, p < 0.001), but this did not quite reach significance for Friction (F_(1, 168)_ = 3.67, p = 0.057). We found no significant main effects in any of the factors for body site or the interaction between age group and body site (all p > 0.05). Figure 1a illustrates the age group differences, where the young group showed significantly higher sensory loadings onto Texture and Moisture than the older group, indicating that they experienced stronger tactile sensations. The third factor, Friction, did not show a significant difference, but shows a similar trend to the other factors. The fourth factor, Temperature, gave positive loadings for warmth and negative for cool, where younger participants perceived self-touch as cool and the older group as warm.

**Figure 1.**
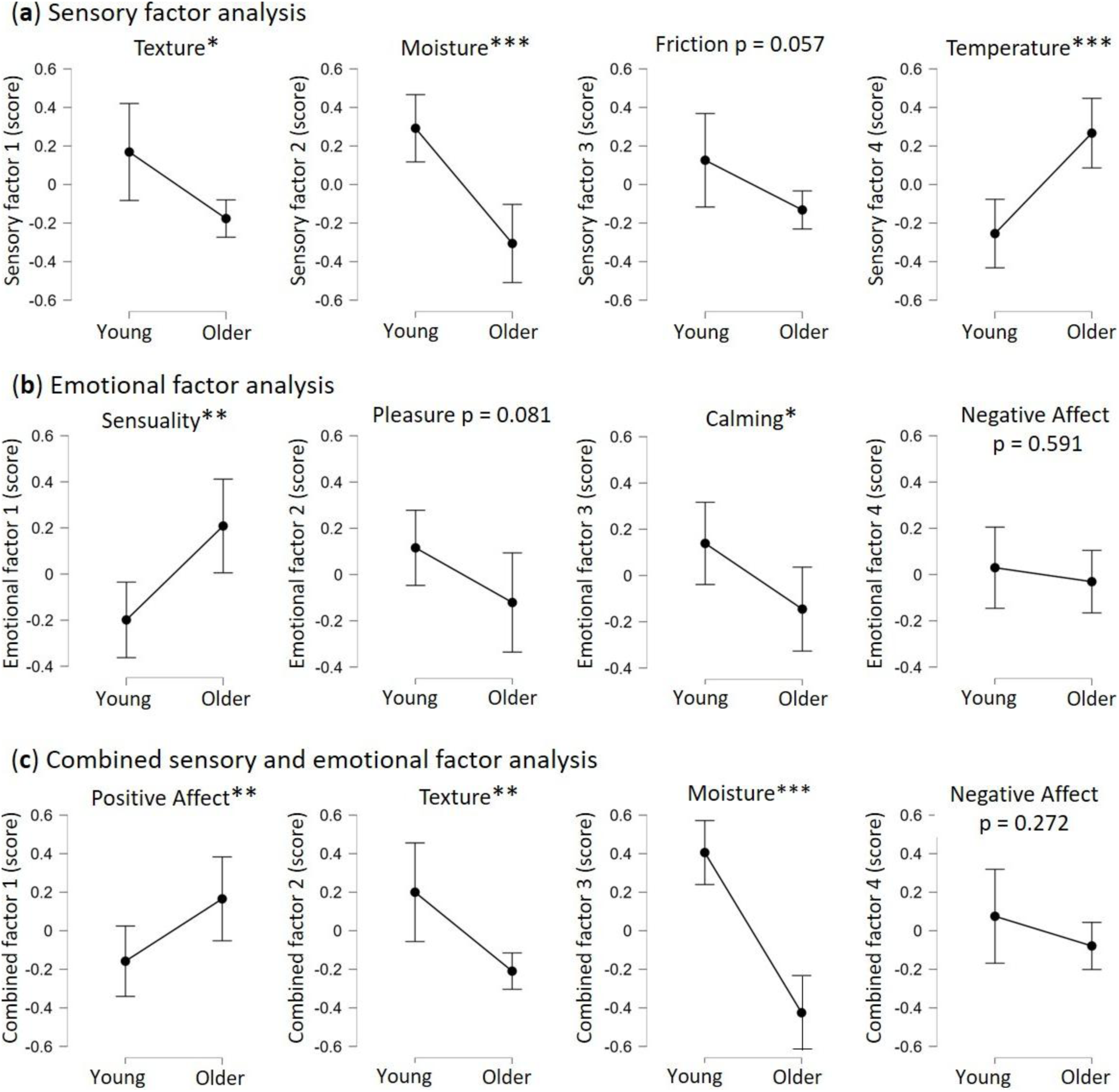
Mean regression output scores for each factor in the three factor analyses per age group. (a) For the sensory factor analysis, four factors were found, where all of the factors, apart from Friction (p = 0.057), showed a significant difference between the young and older age groups. (b) For the emotional factor analysis, four emotional factors were found, where Sensuality and Calming showed a significant difference between the young and older age groups. (c) For the combined sensory and emotional factor analysis, four factors were also found, where all of the factors, apart from Negative Affect, showed a significant difference between the young and older age groups. Significance: *p < 0.05, **p < 0.01, ***p < 0.001. Error bars represent 95% confidence intervals.

**Table 2:** Factor loadings and variance explained by the sensory descriptors factor analysis. We found four factors in the sensory descriptors factor analysis, named Texture, Moisture, Friction, and Temperature. Only descriptors with significant loadings (above 0.3) were included in each factor and the table order indicates their relative contributions.

| Sensory factor 1:<br>Texture<br>15% of variance |  | Sensory factor 2:<br>Moisture<br>12% of variance |  | Sensory factor 3:<br>Friction<br>11% of variance |  | Sensory factor 4:<br>Temperature<br>6% of variance |  |
| --- | --- | --- | --- | --- | --- | --- | --- |
| Sensory descriptor | Regression coefficient | Sensory descriptor | Regression coefficient | Sensory descriptor | Regression coefficient | Sensory descriptor | Regression coefficient |
| Rough | 0.73 | Smooth | 0.74 | Burning | 0.69 | Warm | 0.67 |
| Hairy | 0.70 | Damp | 0.68 | Lumpy | 0.67 | Cold | -0.56 |
| Hard | 0.70 | Slippery | 0.66 | Hot | 0.60 | Damp | -0.50 |
| Gritty | 0.69 | Soft | 0.65 | Bumpy | 0.54 | Vibrating | 0.43 |
| Bumpy | 0.60 | Fluffy | 0.56 | Rubbery | 0.51 | Hot | 0.30 |
| Jagged | 0.53 | Wet | 0.56 | Gritty | 0.47 |  |  |
| Prickly | 0.50 | Cold | 0.44 | Jagged | 0.42 |  |  |
| Firm | 0.41 | Greasy | 0.35 | Dry | 0.35 |  |  |
| Dry | 0.39 | Sticky | 0.31 | Greasy | 0.35 |  |  |
| Fuzzy | 0.39 |  |  | Warm | 0.34 |  |  |
| Lumpy | 0.36 |  |  |  |  |  |  |
| Sharp | 0.36 |  |  |  |  |  |  |

For the emotional factor analysis, four dimensions were found: Sensuality, Pleasure, Calming, and Negative Affect. Table 3 shows the variance explained per factor and the regression loadings. Using ANOVA, we found a significant main effect of age group for Sensuality (F_(1, 168)_ = 9.67, p = 0.002) and Calming (F_(1, 168)_ = 4.93, p = 0.028), whereas the effect was not significant for Pleasure (F_(1, 168)_ = 3.07, p = 0.081) or Negative Affect (F_(1, 168)_ = 0.29, p = 0.591). The older group therefore found self-touch to be a more sensual experience than the young group, whereas the young group found it to be more calming (Figure 1b). We found no significant main effects in any of the factors for body site or the interaction between age group and body site (all p > 0.05).

**Table 3:** Factor loadings and variance explained by the emotional factor analysis. We found four factors in the emotional descriptors factor analysis, Sensuality, Pleasure, Calming, and Negative Affect. Only descriptors with significant loadings (above 0.3) were included in each factor and the table order indicates their relative contributions.

| Emotional factor 1:<br>Sensuality<br>18% of variance |  | Emotional factor 2:<br>Pleasure<br>15% of variance |  | Emotional factor 3:<br>Calming<br>15% of variance |  | Emotional factor 4:<br>Negative Affect<br>6% of variance |  |
| --- | --- | --- | --- | --- | --- | --- | --- |
| Emotional descriptor | Regression coefficient | Emotional descriptor | Regression coefficient | Emotional descriptor | Regression coefficient | Emotional descriptor | Regression coefficient |
| Sensual | 0.72 | Pleasurable | 0.80 | Calming | 0.70 | Irritating | 0.57 |
| Sexy | 0.72 | Comfortable | 0.73 | Soothing | 0.69 | Discomfort | 0.53 |
| Exciting | 0.69 | Enjoyable | 0.72 | Relaxing | 0.61 | Arousing | 0.32 |
| Thrilling | 0.53 | Soothing | 0.34 | Arousing | 0.47 |  |  |
| Desirable | 0.49 | Desirable | 0.32 | Comfortable | 0.37 |  |  |
| Relaxing | 0.37 |  |  | Thrilling | 0.34 |  |  |
| Soothing | 0.32 |  |  |  |  |  |  |

For the combined factorial analyses, where we included all sensory and emotional descriptors together, four dimensions were found: Positive Affect, Texture, Moisture, and Negative Affect. Table 4 shows the variance explained per factor and the regression loadings, where the sensory and emotional descriptors are highlighted in blue and red, respectively. Using ANOVA, we found a significant main effect of age group for Positive Affect (F_(1, 168)_ = 5.14, p = 0.025), Texture (F_(1, 168)_ = 8.51, p = 0.004), Moisture (F_(1, 168)_ = 42.03, p < 0.001), although not for Negative Affect (F_(1, 168)_ = 1.22, p = 0.272). We found no significant main effects in any of the factors for body site or the interaction between age group and body site for this combined analysis (all p > 0.05). Figure 1c illustrates the age group differences, where it can be seen that the older group had significantly higher loadings on Positive Affect than the young group, as per the emotional factor analysis. Similarly, the young group had significantly higher loadings for the Texture and Moisture factors, as we found in the sensory factor analysis. Also like the emotional factor analysis, we found a Negative Affect factor and this was, again, not significant between the groups.

**Table 4:** Factor loadings and variance explained by the combined sensory and emotional descriptors factor analysis. We found four factors in the combined sensory and emotional descriptors factor analysis, Positive Affect, Texture, Moisture, and Negative Affect. Only descriptors with significant loadings (above 0.3) were included in each factor and the table order indicates their relative contributions. All factors had significant loadings of both sensory and emotional words.

| Touch factor 1:<br>Positive affect<br>13% of variance |  | Touch factor 2:<br>Texture<br>11% of variance |  | Touch factor 3:<br>Moisture<br>9% of variance |  | Touch factor 4:<br>Negative Affect<br>9% of variance |  |
| --- | --- | --- | --- | --- | --- | --- | --- |
| Touch descriptor | Regression coefficient | Touch descriptor | Regression coefficient | Touch descriptor | Regression coefficient | Touch descriptor | Regression coefficient |
| Sensual | 0.70 | Rough | 0.73 | Damp | 0.78 | Burning | 0.68 |
| Soothing | 0.69 | Hairy | 0.73 | Smooth | 0.59 | Irritating | 0.61 |
| Relaxing | 0.65 | Gritty | 0.72 | Soft | 0.58 | Hot | 0.58 |
| Desirable | 0.64 | Hard | 0.67 | Cold | 0.57 | Lumpy | 0.55 |
| Exciting | 0.64 | Bumpy | 0.62 | Slippery | 0.62 | Bumpy | 0.52 |
| Sexy | 0.59 | Jagged | 0.58 | Wet | 0.58 | Discomfort | 0.51 |
| Enjoyable | 0.58 | Prickly | 0.49 | Fluffy | 0.45 | Rubbery | 0.50 |
| Thrilling | 0.58 | Lumpy | 0.42 | Pleasurable | 0.43 | Gritty | 0.43 |
| Comfortable | 0.56 | Fuzzy | 0.37 | Comfortable | 0.42 | Warm | 0.40 |
| Pleasurable | 0.54 | Sharp | 0.37 | Enjoyable | 0.41 | Dry | 0.36 |
| Calming | 0.46 | Dry | 0.36 | Greasy | 0.36 | Greasy | 0.35 |
| Warm | 0.34 | Firm | 0.35 | Soothing | 0.33 |  |  |
| Arousing | 0.41 | Arousing | 0.34 | Calming | 0.32 |  |  |
| Soft | 0.38 | Smooth | -0.32 |  |  |  |  |
|  |  | Soft | -0.32 |  |  |  |  |

### Network Analysis

To explore the relationships between descriptors, we conducted network analyses where descriptors are ‘nodes’ and the connection strength between them can be visualized. The whole network can be seen in Figure 2, where the mean weight of the estimated network was 0.15 (numbers of nodes = 40; number of non-zero edges 141/780; sparsity = 0.21), which indicates a moderately-connected network. As a first step including all the data together, the network analysis revealed important insights into how the sensory and emotional descriptors are linked. The majority of connections (edges) between nodes were positive (shown in blue) and notable strong node pairs included Soft-Smooth, Damp-Cold, and Burning-Irritating. Only a couple of significant negative correlations (red connections) were found, notably Warm-Cold and Soft-Prickly.

**Figure 2.**
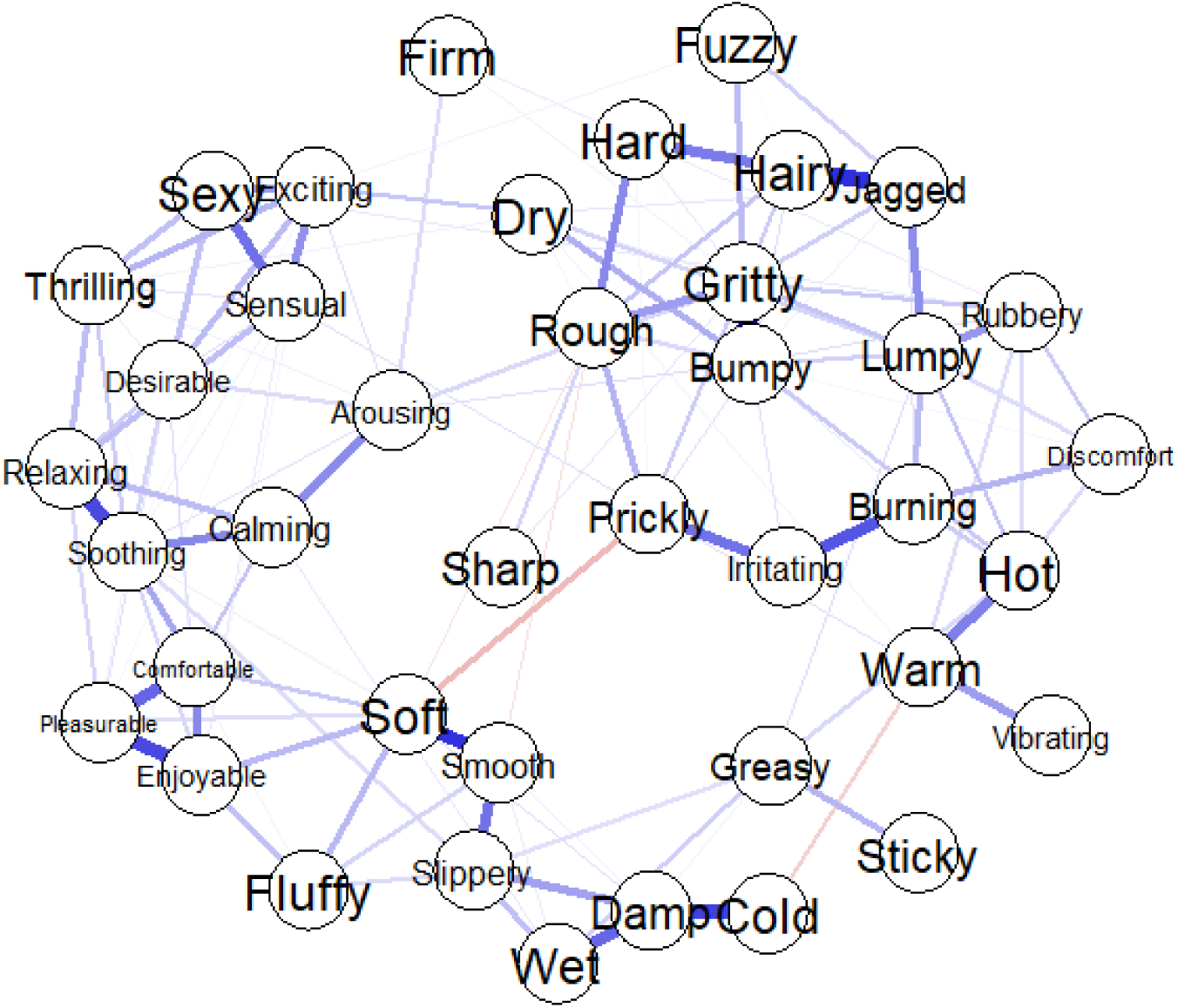
Network analysis of the full dataset to experienced touch. The network visualizes the significant relationships between sensory and emotional descriptors. Blue lines represent positive correlations between nodes, indicating that descriptors are positively associated with each other, while red lines represent negative correlations, showing opposing relationships. Thicker lines indicate stronger correlations.

To analyze the network, we extracted betweenness centrality, which indicates how often a node lies on the shortest path between two other nodes. Nodes with high betweenness act as bridges for connecting clusters of nodes. Significant nodes for betweenness centrality were: Arousing, Hot, Greasy, Gritty, Slippery, Soft, Soothing, and Warm (Fig. 3a). Strength centrality represents the sum of the weights of all edges connected to a node, thus how strongly a node is connected to others in the network. All node descriptors, apart from Discomfort, Firm, Fuzzy, Sharp, Sticky, and Vibrating, showed significant strength centrality (Fig. 3b). Significant nodes play a key role in maintaining the overall structure and connectivity of the network and descriptors including Comfortable, Desirable, Enjoyable, Pleasurable, and Relaxing exhibited smaller confidence intervals, suggesting greater reliability in their centrality. Figure 3c shows significant edges, where green highlights same-type pairs (i.e. sensory-sensory and emotional-emotional), whereas purple shows mixed pairs (i.e. sensory-emotional). There were three significant sensory-emotional pairs: Burning-Irritating, Fluffy-Enjoyable, and Soft-Enjoyable. This shows that the experience of burning was strongly correlated with feelings of irritation, which would be expected given the unpleasant nature of such a sensory input. For Fluffy-Enjoyable, this indicates that light textures were perceived as pleasant, creating positive emotional responses. Similarly, Soft-Enjoyable was significant, also linking softness with an appreciation of a positive emotional state.

**Figure 3.**
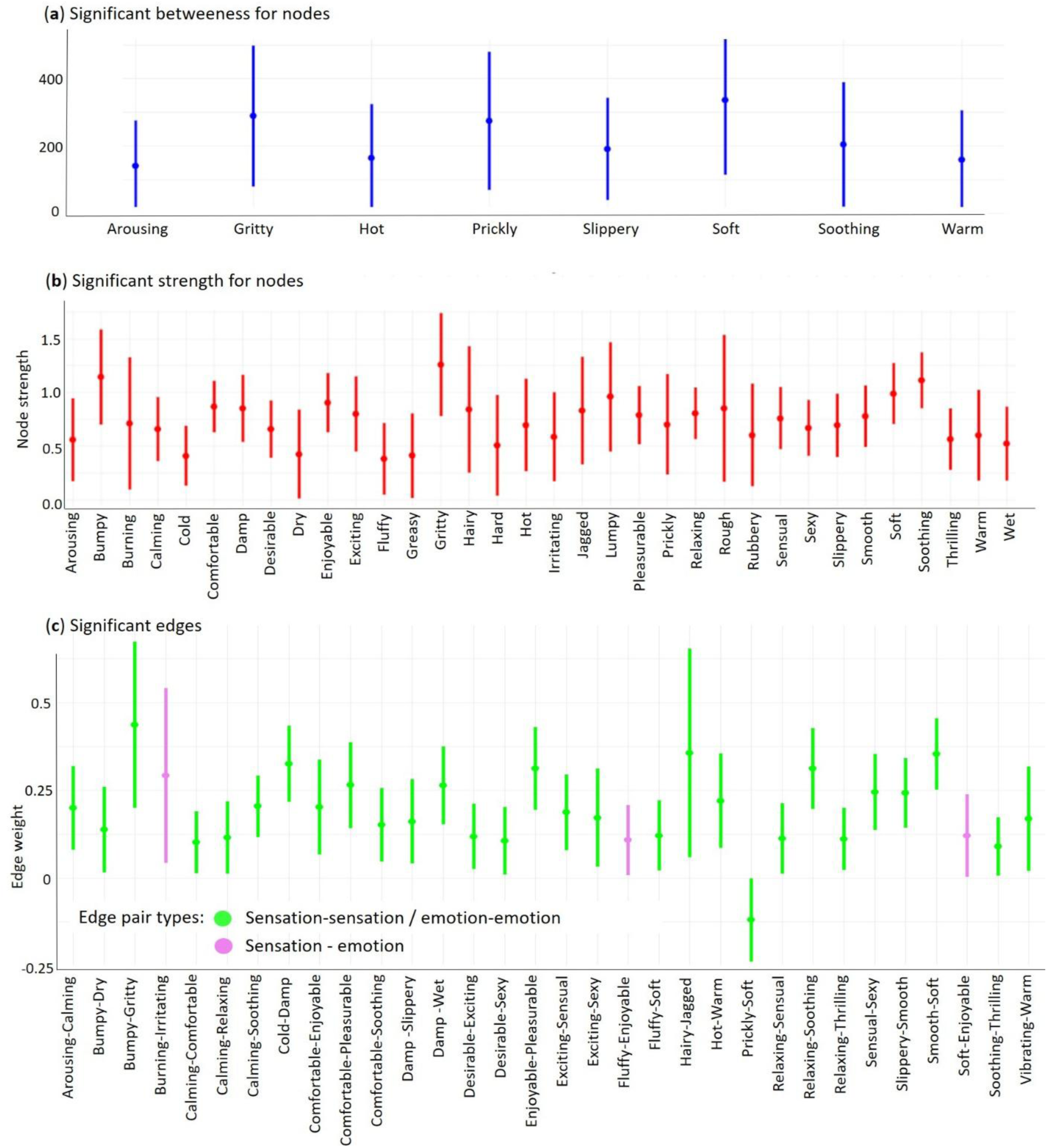
Significant nodes and edges in the full network analysis. Significant nodes (descriptors) and edges (significant relationships between descriptors) in the full network analysis. Significant nodes are shown in (a) for betweenness (in blue) and in (b) for strength centrality (in red). (c) Significant edges, where green represents sensation-sensation and emotion-emotion (i.e. matched) pairings, whereas purple represents mixed sensation-emotion edges. The data are in arbitrary units and all points represent means with ±95% confidence intervals.

We performed a correlation stability analysis to assess the robustness of betweenness and strength using subsampling procedures. This analysis involved progressively omitting participants to examine how network centrality indices maintained their stability. Results indicated that betweenness centrality remained stable (correlation >= 0.7 with original values) when up to 5% of participants were removed, suggesting relatively low robustness. In contrast, strength centrality remained stable even when up to 20% of participants were removed, demonstrating moderate network robustness.

In a second step in the network analysis, we specifically investigated whether age group and body site affected the network. Figure 4 shows the networks for the young and older groups, where it is evident that the older group network has far fewer significant connections. Further, the descriptors Rough and Sharp did not significantly contribute to the network of the older group. To test differences in the networks between age groups, we performed a network comparison test for independent groups which includes a test for global connectivity (measuring overall network density) and a permutation test for clustering coefficients (assessing local clustering patterns). This showed that the overall connectivity in each network was not significantly different (M = 0.512, p = 0.090). However, the permutation test for clustering coefficients differed statistically (difference in global clustering coefficients = 0.566, p < 0.001). This result suggests that there was no difference in node betweenness or strength for the two groups, yet each network had a different structure (i.e. different connections). Local clustering analysis revealed that only one of the young group’s nodes was below a value of 0.75, indicating a highly interconnected structure between the descriptors, whereas nine of the older group’s nodes fell above this value, indicating higher dispersity between descriptors.

**Figure 4.**
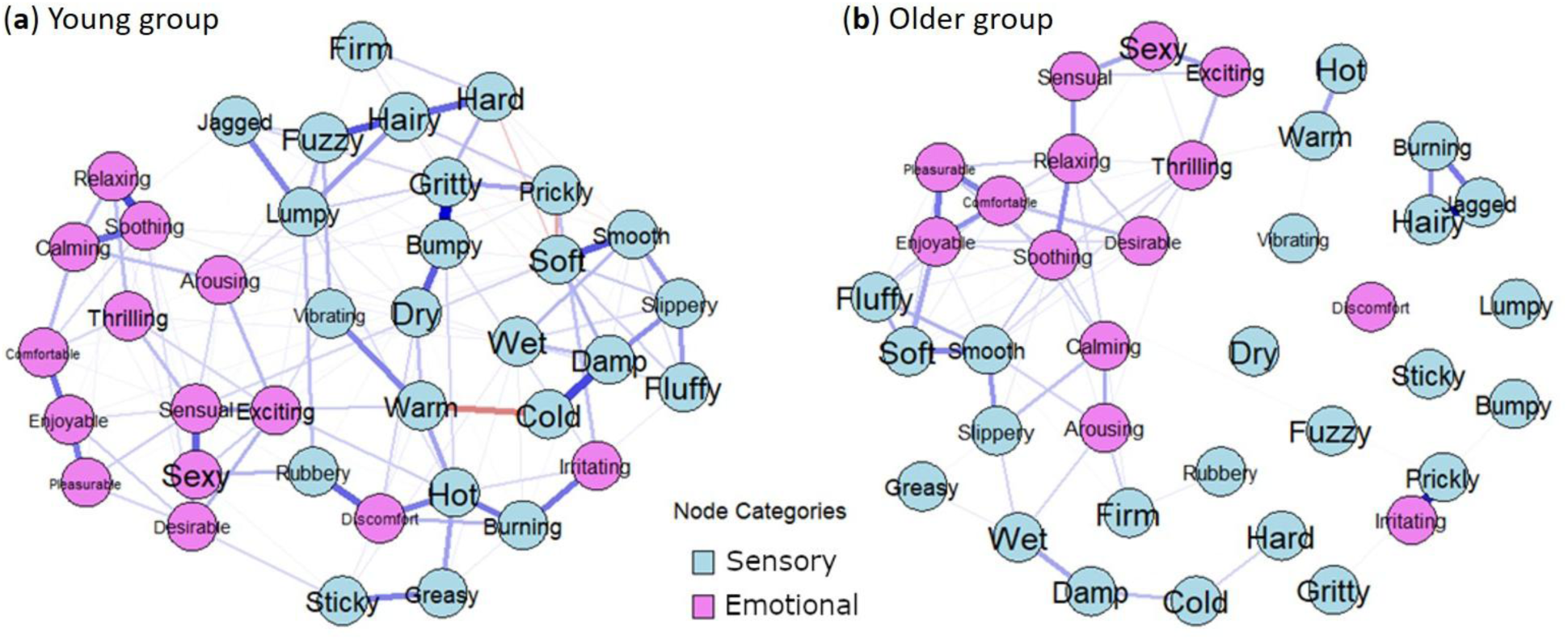
Network analysis of the young and older groups to experienced touch. This network visualization shows the relationships between sensory and emotional descriptors in the (a) young group on the left and (b) older group on the right. Blue lines represent positive correlations and red lines represent negative correlations. Nodes are colored based on their type, with sensory descriptors in light blue and emotional descriptors in pink.

We investigated differences between connections for each age group by examining significant edges (Fig. 5). It can be seen that there are far more significant descriptor pairings (edges) for the young group having a total of 41, as compared to 25 for the older group. However, the older group had 8/25 edges that were mixed sensory-emotional descriptors (pink in Fig. 5), meaning that 32% of the edges were of this mixed type, whereas the young group only had 2/41 (0.05%) edges that were sensory-emotional pairings. Similar to the full network analysis, Burning-Irritating was a significant sensory-emotional edge for the young group and also Hot-Discomfort was found. The older group had very different sensory-emotional pairings, including many positive emotional dimensions (e.g. Smooth-Enjoyable, Soft-Pleasurable, Warm-Comfortable). These differences suggest a stronger link between sensory experiences and positive emotional aspects of touch for the older participants. We conducted a further analysis to test whether there were differences between body sites, performing a network comparison test for dependent groups between the arm and cheek networks. This showed that the overall connectivity was not significantly different (M = 0.295, p = 0.363) and the permutation test was also not significantly different (difference in global clustering coefficients = -0.065, p = 0.722). Overall, this underlines that, like for the factor analysis, there were no significant differences in the perception of touch between these body sites.

**Figure 5.**
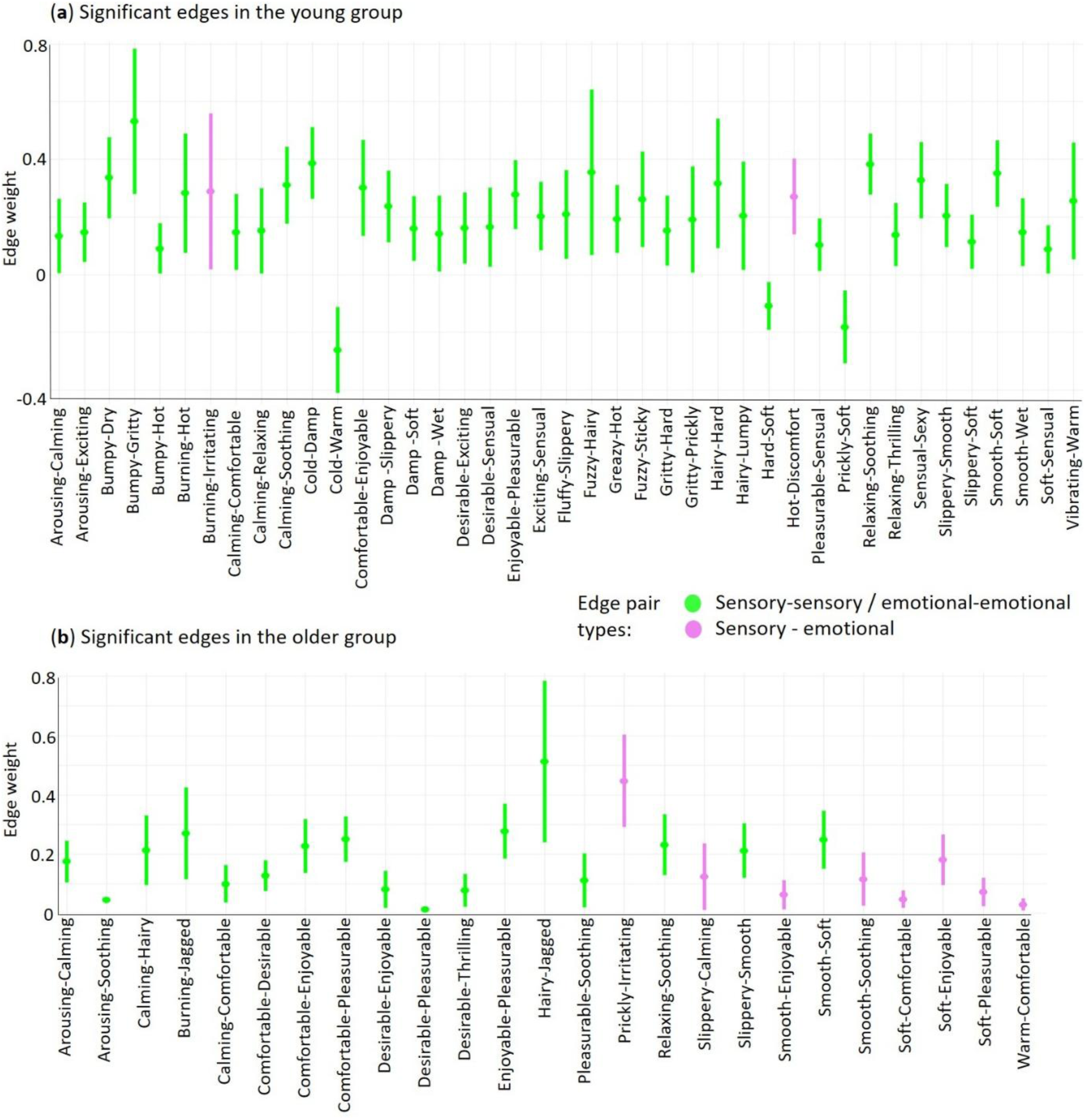
Significant edges in the young and older groups in the network analysis. Significant edges (pairings between node descriptors) and their ±95% confidence intervals for the (a) young group and (b) older group. Green represents sensory-sensory and emotional-emotional (i.e. matched) pairings, whereas purple represents mixed sensory-emotional edges. The data are in arbitrary units and all points represent means with ±95% confidence intervals.

### Clustering analysis

As the permutation test for the age group network analysis resulted in significant differences between the cohesiveness of descriptors, we explored this more deeply through clustering, to visualize the descriptors spatially. This was to validate the permutation test findings for differences between age groups and to uncover latent structures within the multidimensional descriptor space, thereby revealing potential age-related differences in the organization of sensory-emotional representations. Using KMeans clustering on t-SNE axes, descriptors grouped into four clusters for the young group and four clusters for the older group; however, these were different in terms of descriptors per cluster and their spatial dimensions (Figure 6). For the young group, we named the clusters: Arousal, Positive Affect, Texture–Negative Affect, and Moisture, whereas for the older group, they were: Arousal-Moisture, Positive Affect, Negative Affect, and Texture. The silhouette score, which evaluates the quality and consistency of cluster assignments, was 0.60 for the young group and 0.58 for the older group, indicating reasonably well-defined clustering structures, with good separation between clusters and high cohesion within them. All clusters contained both sensory and emotional descriptors, apart from Moisture in the young group and Texture in the older group, which were composed only of sensory descriptors. Although there were similarities in clusters between the groups (e.g. Positive Affect for both), some descriptors moved locations, giving different combinations (e.g. Texture-Negative Affect and Arousal were separate in the young group, whereas, Arousal-Moisture was grouped together, yet Texture and Negative Affect were separate for the older group).

**Figure 6.**
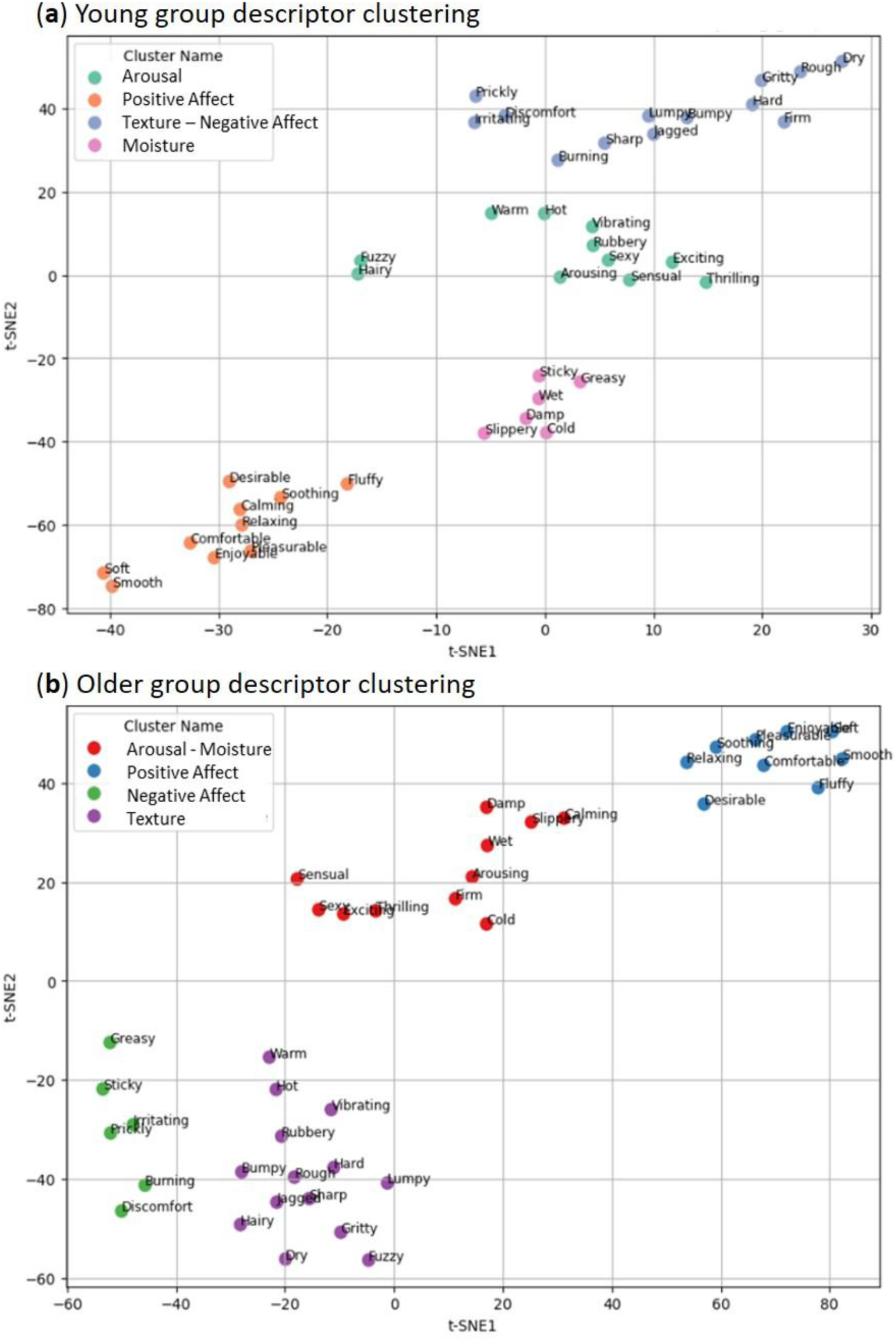
t-SNE plots for descriptor clustering per age group. Plots showing t-distributed Stochastic Neighbor Embedding (t-SNE) clustering results for the (a) young group and (b) older group. Each plot shows that four clusters were found, although these were different between the age groups. The plots provide a two-dimensional representation (X-axis t-SNE1 and Y-axis t-SNE2) of how the descriptors variables relate to each other in lower-dimensional space.

## Discussion

Our study examined the relationships between sensory/discriminative and emotional/affective dimensions in self-touch through the goal-directed grooming task of applying cream to hairy skin, via the hands. Through the French translation of the Touch Perception Task, we found specific interactions between sensory and emotional components, and distinct differences with age. This occurred over different analyses, which we employed to uncover higher dimensions in touch. Firstly, we found that classic factor analysis gave significant differences between age groups, for both sensory and emotional dimensions, apart from no significant difference between age groups for negative affect. Secondly, through novel analyses of the Touch Perception Task, we found that significant descriptors and their connections could be visualized in a network, where there were differences between age groups, but only in the connectivity of the network and not in descriptor localizations. There were far fewer significant descriptor pairings in the older group, although there was a much higher proportion of mixed sensory-emotional pairings, as compared to the young group. Finally, we applied clustering to the networks for each age group and found similar general clusters, but with different components. Therefore, the younger group perceived the sensory aspects of touch more strongly, whereas the older group found the positive affective emotional aspects to be more intense.

Our complementary statistical approaches demonstrate the profound effects of aging on tactile perception, extending previous work showing that tactile sensitivity reduces [10,52], and that touch becomes more pleasant [28,32,33], with age. We found systematic decreases in sensory tactile perception in the older group, such as for texture and moisture. This implies a diminished capacity for detecting finer aspects of touch, as would be expected with peripheral and/or central tactile decline, especially at the hand [9,10,28,52]. In a related investigation on the same participants [28], the older group had significantly drier skin on their finger, but not on the arm or cheek, demonstrating further alterations in skin anatomy and physiology with age [53]. These changes in touch at the finger therefore have clear consequences for self-touch perception. For emotional tactile aging, previous work has measured pleasantness ratings [28,33], yet we captured emotional/affective touch in a broader way, including negative aspects. We add to growing evidence that touch becomes more positive with age, while also finding that negative affective touch (e.g. discomfort) remains stable. However, we did not include conditions using particularly negative stimuli (e.g. sticky liquids, [54]), therefore the impact of negative affective touch should be further explored. The origin of the increase of positive affect remains unknown. Sehlstedt et al. [33] compared pleasantness in touch and odors, showing that only tactile pleasantness increased with age, meaning that a general appreciation of sensory input with age seems unlikely. The increase in positive tactile affect could stem from peripheral and/or central mechanisms [55]. Decreases in peripheral innervation with age could change the balance between fiber classes, but the specific proportion of loss for different fiber types, such as CT afferents, remains unknown. Centrally, aging has an impact on tactile processing that can be accompanied by cognitive decline, where older adults rely more on emotional mechanisms [55]. This is in line with the increase in sensory-emotional descriptor pairings that we found in the network analysis for the older group.

Although aging effects were evident, we found no effect of body site touched in any of our analyses, although this was not unexpected, as both the cheek and forearm are hairy skin sites that are both highly involved in affective touch. An earlier study found that passive stroking touch to the cheek was rated as more pleasant than to the forearm, although both were very pleasant [56]. A later study using the Touch Perception Task found no differences between touch to the forearm and cheek for any sensory or emotional factor, although found that active touch using textures was less pleasant overall than being passively touched with these [1]. These differences may therefore be due to the mode of touch, where touch perception is inherently different in active, self-directed touch [23,24]. However, this is important to consider in self-touch, such as in self-care and grooming, where the hand may play a critical role in shaping touch to one’s own body [25,28].

We had a secondary objective to apply advanced analytical methods to the Touch Perception Task, as newer statistical approaches could be applied to map how tactile dimensions evolve with each other. As a control, our first approach followed the classic factor analysis method set-out in the original Touch Perception Task [2], where sensory and emotional descriptors were dealt with separately, as the emotional dimensions of semantic-perceptual space had previously been unexplored. Guest et al. [2] did briefly investigate relationships between sensory and emotional factors, but this was not explored in-depth, as the authors explicitly chose to study the emotional side of touch separately and compare it to sensory dimensions that had been studied in other work [57,58]. The Touch Perception Task has since been translated into other languages and used in different ways [1,37–41], yet sensory and emotional components have always been considered separately, as per the original study [2]. Therefore, we went beyond this to relate these two aspects in touch in a combined factor analysis, especially as recent papers emphasize the relationships between sensory/discriminative and emotional/affective touch and that they occur together, even possibly being inseparable [3,59,60].

When we analyzed the sensory and emotional descriptors separately, each gave four significant factors, but when analyzed together, four significant factors were also produced, which all included some sensory and emotional descriptors; however, each factor was more biased to one side or the other. For example, the first combined factor was Positive Affect that also included the sensory descriptors Warm and Soft. The second factor, Texture, was mainly sensory, but included Arousing. The third factor, Moisture, was relatively mixed between sensory and emotional descriptors, with a sensory dominance. The final factor, termed Negative Affect, was composed mainly of sensory descriptors, but included Irritating and Discomfort, demonstrating that sensory aspects contribute to negative tactile experiences. This combined analysis showed that even though there are fewer emotional descriptors in the Touch Perception Task, the emotional aspect is very present and even dominates as the primary factor. These findings motivated the further exploration of how all the descriptors related to each other, applying different statistical approaches to the data.

The network analysis revealed the complexities of how the descriptors were positioned in space, their dominance over the network, and the strength of how they corresponded to each other, allowing us to visualize and quantify these relationships. The vast majority of significant connections between descriptors were positive, although some were negative, such as Warm-Cold and Soft-Prickly. However, in the older group, there were far fewer connections between nodes, especially for sensory descriptors, and none of these were negative. The clustering analysis showed differences in the spatial positioning of the descriptors for the age groups, where there was a strong Positive Affect cluster for both groups, which was tightly-clustered in the older group, but that the other sensory and Negative Affect clusters shifted in terms of descriptor content and had more distant spatial relations with age.

The present work represents an advance in the way holistic tactile experiences can be interpreted, yet there are limitations. Firstly, only women were studied and there could be differences in tactile perception depending on sex. A large meta-analysis found that women had slightly increased tactile pleasantness ratings over men [61], yet this was a small effect and it is unknown how this evolves with age. We also only tested touch to two body sites, the cheek and forearm, thus more sites could be investigated in the future, to build body maps for higher-level tactile dimensions. Further, we used skin-to-skin self-touch in the application of a cream, whereas the perception of touch may change if self-touch is applied via a tool, as used in previous work [1,2,32]. Finally, although we tested over 40 participants and our various analyses produced highly convergent results, more data are needed to develop robust models of tactile experiences. Thus, we present a path towards the modeling of complex touch perception data that could then be used to make predictions about relationships in tactile dimensions, such as how to increase or decrease emotional aspects via the manipulation of sensory aspects. Classically, linear statistics are used to interpret touch perception data, yet our clustering allowed a different visualization in a non-linear, less-constrained way. Both the network and clustering analyses provide new bases for further exploring tactile dimensions, where machine learning approaches could be applied [62,63]. These could capture numerous aspects in touch, thus not only sensory and emotional aspects, but also differences with age, body site, texture touching the skin, and mode of touch.

In conclusion, our study contributes to growing evidence demonstrating the complex interplay between sensory and emotional tactile percepts and their changes with age. As well as providing a French translation of the Touch Perception Task, we find that while sensory/discriminative tactile dimensions diminish with age, positive emotional/affective dimensions strengthen. This impacts the way in which we interact with ourselves and others, as well as the design of products, where considering the emotional aspects is more important, especially for older people [64,65]. Our findings underscore the limitations of classic statistical methods in capturing the complexity of the full tactile experience, where these conventional approaches may fail to account for the intricate, non-linear interactions between descriptors, as found in the nuanced patterns observed. Thus, there is a need to employ advanced, multidimensional methods, to unravel the complexity of tactile perception more effectively, creating a framework to interpret touch. Future work could take this further by applying techniques such as deep learning and the prediction of causal relationships in tactile experiences.

## Acknowledgements

The work was supported by an Agence National de la Recherche (ANR, France) grant ‘UNTOUCH’ no. ANR-22-ORAR-0007 to R.A. in the Open Research Area (ORA7) collaboration call. L’Oréal provided funding to conduct the experiments.

## Author contributions

MRB, JMA, RJ, and RA conceptualized the work. MD performed the experiments, with help from RA and JMA. RA performed pre-processing of the data and initial analyses. MRB performed the analyses with the help of PB, with guidance from RA and RJ. MRB drafted the first version of the manuscript. MRB, PB, AC, RJ, and RA worked on the writing and interpretation. All authors contributed to the work and approved the final version. All authors agree to be accountable for all aspects of the work in ensuring that questions related to the accuracy or integrity of any part of the work are appropriately investigated and resolved.

## Data availability statement

The data are available in the OSF repository https://osf.io/h9k37.

## References

1. Ackerley, R., Saar, K., McGlone, F. & Backlund Wasling, H. Quantifying the sensory and emotional perception of touch: differences between glabrous and hairy skin. Frontiers in Behavioral Neuroscience 8, 34 (2014).

2. Guest, S. et al. The development and validation of sensory and emotional scales of touch perception. *Attention*, Perception & Psychophysics 73, 531–550 (2011).

3. Schirmer, A., Croy, I. & Ackerley, R. What are C-tactile afferents and how do they relate to “affective touch”? Neuroscience & Biobehavioral Reviews 151, 105236 (2023).

4. Perini, I., Olausson, H. & Morrison, I. Seeking pleasant touch: neural correlates of behavioral preferences for skin stroking. Frontiers in Behavioral Neuroscience 9, 8 (2015).

5. McGlone, F., Wessberg, J. & Olausson, H. Discriminative and affective touch: sensing and feeling. Neuron 82, 737–55 (2014).

6. Morrison, I., Löken, L. S. & Olausson, H. The skin as a social organ. Experimental Brain Research 204, 305–14 (2010).

7. van Paasschen, J., Walker, S. C., Phillips, N., Downing, P. E. & Tipper, S. P. The effect of personal grooming on self-perceived body image. International Journal of Cosmetic Science 37, 108–15 (2015).

8. Spille, J. L., Grunwald, M., Martin, S. & Mueller, S. M. Stop touching your face! A systematic review of triggers, characteristics, regulatory functions and neuro-physiology of facial self touch. Neuroscience & Biobehavioral Reviews 128, 102–116 (2021).

9. Corniani, G. & Saal, H. P. Tactile innervation densities across the whole body. Journal of Neurophysiology 124, 1229–1240 (2020).

10. Stevens, J. C. & Choo, K. K. Spatial Acuity of the Body Surface over the Life Span. Somatosensory & Motor Research 13, 153–166 (1996).

11. Weinstein, S. Intensive and Extensive Aspects of Tactile Sensitivity as a Function of Body Part, Sex, and Laterality. in The Skin Senses (ed. Kenshalo, D.) 195–222 (Charles C. Thomas, Springfield, IL, 1968).

12. Vallbo, Å., Olausson, H. & Wessberg, J. Unmyelinated afferents constitute a second system coding tactile stimuli of the human hairy skin. Journal of Neurophysiology 81, 2753–2763 (1999).

13. Nordin, M. Low-threshold mechanoreceptive and nociceptive units with unmyelinated (C) fibres in the human supraorbital nerve. Journal of Physiology 426, 229–240 (1990).

14. Watkins, R. H. et al. Evidence for sparse C-tactile afferent innervation of glabrous human hand skin. Journal of Neurophysiology 125, 232–237 (2021).

15. Löken, L. S., Wasling, H. B., Olausson, H., McGlone, F. & Wessberg, J. A topographical and physiological exploration of C-tactile afferents and their response to menthol and histamine. Journal of Neurophysiology 127, 463–473 (2022).

16. Ackerley, R. et al. Human C-tactile afferents are tuned to the temperature of a skin-stroking caress. Journal of Neuroscience 34, 2879–83 (2014).

17. Löken, L. S., Wessberg, J., Morrison, I., McGlone, F. & Olausson, H. Coding of pleasant touch by unmyelinated afferents in humans. Nature Neuroscience 12, 547–8 (2009).

18. Ackerley, R., Carlsson, I., Wester, H., Olausson, H. & Backlund Wasling, H. Touch perceptions across skin sites: differences between sensitivity, direction discrimination and pleasantness. Frontiers in Behavioral Neuroscience 8, 54 (2014).

19. Stevens, J. C. & Choo, K. K. Temperature sensitivity of the body surface over the life span. Somatosensory & Motor Research 15, 13–28 (1998).

20. Valenza, A., Bianco, A. & Filingeri, D. Thermosensory mapping of skin wetness sensitivity across the body of young males and females at rest and following maximal incremental running. Journal of Physiology 597, 3315–3332 (2019).

21. Nummenmaa, L., Glerean, E., Hari, R. & Hietanen, J. K. Bodily maps of emotions. Proceedings of the National Academy of Sciences of the United States of America 111, 646–51 (2014).

22. Fagard, J., Esseily, R., Jacquey, L., O’Regan, K. & Somogyi, E. Fetal Origin of Sensorimotor Behavior. Frontiers in Neurorobotics 12, (2018).

23. Blakemore, S. J., Goodbody, S. J. & Wolpert, D. M. Predicting the consequences of our own actions: the role of sensorimotor context estimation. Journal of Neuroscience 18, 7511–8 (1998).

24. Blakemore, S. J., Wolpert, D. M. & Frith, C. D. Central cancellation of self-produced tickle sensation. Nature Neuroscience 1, 635–40 (1998).

25. Ackerley, R. et al. An fMRI study on cortical responses during active self-touch and passive touch from others. Frontiers in Behavioral Neuroscience 6, 51 (2012).

26. Simões-Franklin, C., Whitaker, T. A. & Newell, F. N. Active and passive touch differentially activate somatosensory cortex in texture perception. Human Brain Mapping 32, 1067–1080 (2011).

27. Cruciani, G., Zanini, L., Russo, V., Boccardi, E. & Spitoni, G. F. Pleasantness ratings in response to affective touch across hairy and glabrous skin: a meta-analysis. Neuroscience & Biobehavioral Reviews 131, 88–95 (2021).

28. Dione, M., Watkins, R. H., Aimonetti, J.-M., Jourdain, R. & Ackerley, R. Effects of skin moisturization on various aspects of touch showing differences with age and skin site. Scientific Reports 13, 17977 (2023).

29. Muret, D. & Dinse, H. R. Tactile learning transfer from the hand to the face but not to the forearm implies a special hand-face relationship. Scientific Reports 8, 11752 (2018).

30. McIntyre, S. et al. The Language of Social Touch Is Intuitive and Quantifiable. Psychological Sciemce 33, 1477–1494 (2022).

31. Samain-Aupic, L., Dione, M., Ribot-Ciscar, E., Ackerley, R. & Aimonetti, J.-M. Relations between tactile sensitivity of the finger, arm, and cheek skin over the lifespan showing decline only on the finger. Frontiers in Aging Neuroscience 16, (2024).

32. Schlintl, C. & Schienle, A. Evaluation of Affective Touch: A Comparison Between Two Groups of Younger and Older Females. Experimental Aging Research 50, 568–576 (2024).

33. Sehlstedt, I. et al. Gentle touch perception across the lifespan. Psychology and Aging 31, 176–184 (2016).

34. Bell-Krotoski, J., Weinstein, S. & Weinstein, C. Testing Sensibility, Including Touch-Pressure, Two-point Discrimination, Point Localization, and Vibration. Journal of Hand Therapy 6, 114–123 (1993).

35. Essick, G. K. et al. Quantitative assessment of pleasant touch. Neuroscience & Biobehavioral Reviews 34, 192–203 (2010).

36. Sailer, U., Hausmann, M. & Croy, I. Pleasantness Only?: How Sensory and Affective Attributes Describe Touch Targeting C-Tactile Fibers. Experimental Psychology 67, 224–236 (2020).

37. Drewing, K., Weyel, C., Celebi, H. & Kaya, D. Systematic Relations between Affective and Sensory Material Dimensions in Touch. IEEE Transactions in Haptics 11, 611–622 (2018).

38. Guest, S. et al. Perception of fluids with diverse rheology applied to the underarm versus forearm skin. Somatosensory & Motor Research 29, 89–102 (2012).

39. Guest, S. et al. Physics and tactile perception of fluid-covered surfaces. Journal of Texture Studies 43, 77–93 (2012).

40. McGlone, F. et al. Touching and feeling: differences in pleasant touch processing between glabrous and hairy skin in humans. European Journal of Neuroscience 35, 1782–8 (2012).

41. van Hooijdonk, R. et al. Touch-induced pupil size reflects stimulus intensity, not subjective pleasantness. Experimental Brain Research 237, 201–210 (2019).

42. Kaiser, H. F. An index of factorial simplicity. Psychometrika 39, 31–36 (1974).

43. Friedman, J., Hastie, T. & Tibshirani, R. Sparse inverse covariance estimation with the graphical lasso. Biostatistics 9, 432–441 (2008).

44. Hittner, J. B., May, K. & Silver, N. C. A Monte Carlo Evaluation of Tests for Comparing Dependent Correlations. The Journal of General Psychology 130, 149–168 (2003).

45. Costantini, G. et al. State of the aRt personality research: A tutorial on network analysis of personality data in R. Journal of Research in Personality 54, 13–29 (2015).

46. Borsboom, D. et al. Network analysis of multivariate data in psychological science. Nature Reviews Methods Primers 1, 58 (2021).

47. Epskamp, S., Waldorp, L. J., Mõttus, R. & Borsboom, D. The Gaussian Graphical Model in Cross-Sectional and Time-Series Data. Multivariate Behavioral Research 53, 453–480 (2018).

48. Van Borkulo, C. D. et al. Comparing network structures on three aspects: A permutation test. Psychological Methods 28, 1273–1285 (2023).

49. Epskamp, S., Borsboom, D. & Fried, E. I. Estimating Psychological Networks and their Accuracy: A Tutorial Paper. Preprint at http://arxiv.org/abs/1604.08462 (2017).

50. Friedman, B. H. et al. Autonomic characteristics of nonclinical panic and blood phobia. Biological Psychiatry 34, 298–310 (1993).

51. Friedman, J., Hastie, T. & Tibshirani, R. Sparse inverse covariance estimation with the graphical lasso. Biostatistics 9, 432–441 (2008).

52. Samain-Aupic, L., Dione, M., Ribot-Ciscar, E., Ackerley, R. & Aimonetti, J.-M. Relations between tactile sensitivity of the finger, arm, and cheek skin over the lifespan showing decline only on the finger. Open Science Framework 10.17605/OSF.IO/879KB (2024) doi:10.17605/OSF.IO/879KB.

53. Ezure, T., Amano, S. & Matsuzaki, K. Aging-related shift of eccrine sweat glands toward the skin surface due to tangling and rotation of the secretory ducts revealed by digital 3D skin reconstruction. Skin Research and Technology 27, 569–575 (2021).

54. Cavdan, M., Fehlberg, M., Bennewitz, R. & Drewing, K. Gooey stuff: the psychophysics of unpleasantness in response to touching liquids. Royal Society Proceedings B 292, 20252244 (2025).

55. McIntyre, S., Nagi, S. S., McGlone, F. & Olausson, H. The Effects of Ageing on Tactile Function in Humans. Neuroscience 464, 53–58 (2021).

56. Essick, G. K., James, A. & McGlone, F. P. Psychophysical assessment of the affective components of non-painful touch. Neuroreport 10, 2083–7 (1999).

57. Hollins, M., Faldowski, R., Rao, S. & Young, F. Perceptual dimensions of tactile surface texture: a multidimensional scaling analysis. Perception & Psychophysics 54, 697–705 (1993).

58. Hollins, M., Bensmaïa, S., Karlof, K. & Young, F. Individual differences in perceptual space for tactile textures: evidence from multidimensional scaling. Perception & Psychophysics 62, 1534–1544 (2000).

59. Morrison, I. Affective and discriminative touch: A reappraisal. Current Opinion in Behavioral Sciences 43, 145–151 (2022).

60. Schirmer, A., Lai, O., McGlone, F., Cham, C. & Lau, D. Gentle stroking elicits somatosensory ERP that differentiates between hairy and glabrous skin. *Social*, Cognitive & Affective Neuroscience 17, 864–875 (2022).

61. Russo, V., Ottaviani, C. & Spitoni, G. F. Affective touch: A meta-analysis on sex differences. Neuroscience & Biobehavioral Reviews 108, 445–452 (2020).

62. Contractor, N. S., Wasserman, S. & Faust, K. Testing Multitheoretical, Multilevel Hypotheses About Organizational Networks: An Analytic Framework and Empirical Example. AMR 31, 681– 703 (2006).

63. Rubinov, M. & Sporns, O. Complex network measures of brain connectivity: Uses and interpretations. NeuroImage 52, 1059–1069 (2010).

64. Roso, A. et al. Contribution of cosmetic ingredients and skin care textures to emotions. International Journal of Cosmetic Science 46, 262–283 (2024).

65. Sato, C. Changes in Emotions and Attitudes When Using Cosmetics. Journal of the Society for Cosmetic Chemists Japan 54, 351–357 (2020).

