## Supplementary Figure 1 for "Changes of sensory and emotional tactile dimensions with age in self-touch via the application of creams"

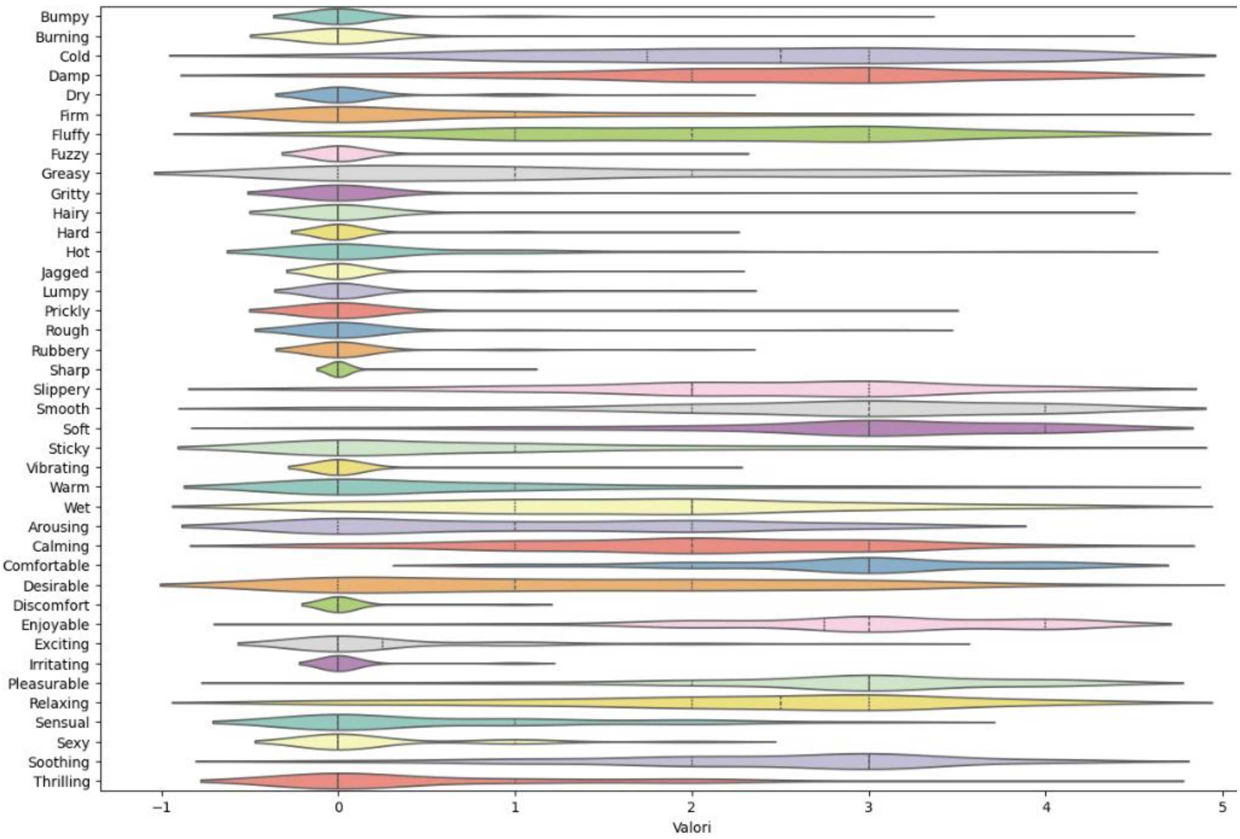


**Supplementary Figure 1. Violin plot of the distribution of descriptor ratings.**

Each descriptor is represented on the y-axis, with its corresponding rating value as a violin plot on the x-axis. The width of each violin indicates the density of the data at different values, giving a sense of the distribution's spread. The central line within each violin marks the median, to compare the central tendency between descriptors.
